# The Emerging Gait Dysfunction Phenotype in Idiopathic Parkinson’s Disease

**DOI:** 10.1101/632638

**Authors:** Frank M. Skidmore, William S Monroe, Christopher Hurt, Anthony P Nicholas, Adam Gerstenecker, Thomas Anthony, Leon Jololian, Gary Cutter, Adil Bashir, Thomas Denny, David Standaert, Elizabeth A Disbrow

## Abstract

**Objective:** Severity of motor symptoms in Parkinson’s disease (PD), and rate of change of these symptoms, suggests the existence of disease subgroups. One important PD subgroup is defined by postural instability and gait dysfunction (PIGD), which is associated with disability, lower quality of life, and cognitive deterioration. In this study, we evaluate what clinical factors at baseline are associated with early development of postural instability in PD.

**Methods:** Data was downloaded from the Parkinson’s Progressive Markers Initiative (PPMI). Several clinical features predict development of postural instability. We provisionally term the associated phenotype the emerging gait disorder (eGD) phenotype. We evaluate validity of the phenotype in two held-out populations.

**Results:** Individuals with the proposed eGD phenotype have a significantly higher risk of developing postural instability in both validation sets (p < 0.00001 in both sets). The proposed eGD phenotype occurred before development of postural instability (HY stage ≥ 3) in 289 of 301 paired comparisons (Fischer Exact Test, p < 0.000001), with a median progression time from development of eGD phenotype to postural instability of 972 days. Individuals with the proposed eGD phenotype at baseline had more rapid cognitive decline as measured by the Montreal Cognitive Assessment (p = 0.002) and Hopkins Verbal Learning Test (Total Recall, p = 0.008).

**Interpretation:** We describe a clinical phenotype, detectable at baseline in a subset of individuals with PD, that is associated with accelerated development of postural instability. Within the sample, development of the eGD phenotype reliably precedes development of disability, and is a harbinger of more rapid cognitive progression.

## INTRODUCTION

Severity of motor symptoms in PD, and rate of change of these symptoms over time is quite variable, suggesting the existence of disease subgroups with varying rates of progression.^1-5^ Amongst this variability, individuals with PD universally share symptomatic dopamine deficiency, which has led to a wave of innovation to assist with management of dopaminergic-related symptoms. However, treating PD as a monomorphic condition has, as noted by Espay et al., “consistently failed when testing potential disease-modifying interventions.^6^

One long recognized transition point in the development of PD-related disability is the development of postural instability. The relationship between gait and balance and disability was first examined through the prism of the Hoehn and Yahr (HY)^7,8^ scale, the first widely used PD severity scale (Table 1: Recapitulation of HY Scale). Jankovic et al. later developed a postural instability/gait dysfunction (PIGD)^9-11^ score, derived from the Unified Parkinson’s Disease Rating Scale (UPDRS),^12,13^ a structured history and clinical exam. Development of the PIGD phenotype has been associated with disability and lower quality of life,^14-16^ more rapid progression of both cognitive and motor dysfunction,^17-19^ and substantially higher risk of developing diffuse lewy body disease (DLB) and Parkinson’s Disease Dementia (PDD).^20,21^ Predictably, advancing HY status is similarly associated with reduced quality of life and cognitive decline.^22,23^ However, in the absence of other disabilities individuals with PD are HY stage 1 or 2 at onset of disease. Similarly, the PD PIGD phenotype is based on clinical parameters in later stages of disease,^10-12^ but is an unstable classification in early disease.^24,25^

**Table 1:** The Hoehn and Yahr Scale^7,8^. The Hoehn and Yahr staging classification predates the more recently employed PIGD formulation and remains in common use. Note: HY stages 3, 4, and 5 describe various degrees of postural instability.

| **Table 1: The Hoehn and Yahr Scale<sup>7,8</sup>** |  |
| --- | --- |
| **Stage** | **Description** |
| 1 | Unilateral involvement usually with minimal or no functional disability |
| 2 | Bilateral or midline involvement without impairment of balance |
| 3 | Bilateral disease: mild to moderate disability with impaired postural reflexes; physically independent |
| 4 | Severely disabling disease; still able to walk or stand unassisted |
| 5 | Confinement to a bed or wheelchair unless aided |

A current, critical need in the field is to develop methods for early identification of individuals at risk of rapid motor progression. Early identification of individuals at risk of progression to postural instability, in particular will serve two important goals. First, as noted above, development of postural instability is associated with global progression of disease, and accordingly identification of at-risk individuals in this context also identifies individuals at higher risk of disability and cognitive progression. Second, selecting individuals with more rapid motor progression will improve the ability to detect the impact of potential disease modifying agents. In particular, while medication has an impact on gait in PD, in the PIGD phenotype gait is less responsive to medication. Clinical trials design in PD is impacted by the problem of disentangling medication response from disease progression.^26,27^ More effective identification of a phenotype (PIGD) with rapid deterioration of a partially medication-unresponsive symptom may mitigate some of these trial design issues, providing the potential of a cleaner signal of presence (or absence) of a disease modifying effect.

We accordingly set out to determine if there are clinical features – a “risk phenotype” – for gait disorder that is detectable at baseline. We developed a risk profile in a derivation set, and evaluated the validity of this measure in two validation sets. To operationalize our findings, we labeled our provisional phenotype the “emerging gait disorder” (eGD) phenotype, and present our results in the form of a clinical scale, the emerging gait disorder screening scale (eGDSS).

## METHODS

### Selection of Sample

Data was downloaded from the Parkinson’s Progressive Markers Initiative (PPMI)^28^ in January of 2019. We screened for data sets that contained at least 5 years of clinical data and identified 380 individuals with de-novo idiopathic PD in PPMI who had at least 5 years of clinical data. A derivation idiopathic PD (dIPD) set of 301, and a validation idiopathic PD (vIPD) set of 79 were developed. The vIPD sample was developed based on availability of imaging in this sample (see Companion Paper, “Imaging Characteristics of the Emerging Gait Phenotype in Idiopathic PD”).^29^ For additional validation, we selected the PPMI genetic cohort, restricting our sample to individuals with HY status of 2 or better (no gait disturbance). PD inclusion criteria in this group were diagnosis of PD for ≤ 7 years, and HY ≤ 4 at entry. All individuals in this sample had mutations in the synuclein alpha (SNCA), leucine-rich repeat kinase 2 (LRRK2), or glucocerebrosidase 1 (GBA1) gene. The genetic PD (GPDv) cohort at the time of this paper contained 220 enrolled individuals. Within the cohort, 141 individuals met inclusion criteria. As a comparative sample, we identified 183 healthy controls who similarly had at least 5 years of clinical follow up. All individuals in PPMI with PD have an “on medication” evaluation at each visit. Thus we study dIPD=301, vIPD=79, vGPD=141, Controls=183. For the purposes of this study, we include exclusively the “on-medicine” evaluation, as we focus specifically on individuals in whom a treatment effect is not adequate to prevent disabling gait related symptoms.

### Sample Features

Data available in PPMI, and included in our analysis, included age and gender. Pertaining to motor assessments, we included the modified Unified Parkinson’s Disease Rating Scale, and HY rating. Given our interest in gait and balance, we calculated PIGD score at each time point over a minimum of 5 years. Cognitive measures included the Montreal Cognitive Assessment (MOCA), the Hopkins Verbal Learning Test (HVLT), Letter Number Sequencing (LNS), Judgement of Line Orientation (JLO), and Symbol-Digit Modality (SDM). All evaluations were performed at baseline and at multiple time points, allowing us to calculate both baseline differences and rate of change by group. Slope of cognitive change in this sample was expressed by change in mean score value divided by time in years.

For **<u>Scale Development and Sample Selection</u>**, Hoehn and Yahr Stage (HY),^7,8^ was dichotomized in the derivation set (301 subjects) to categorize individuals who by year 5 had developed gait dysfunction (Y5_HY3-5) and those who had not (Y5_HY0-2). The HY is a qualitative scale scored from 0-5 (see Table 1). A score of 3 or higher indicates the individual has developed clinically detectable postural instability during stance. We classified a subject as Y5_HY3-5 if at any time over the first 5 years a clinical rating of HY ≥ 3 was scored by any rater while “on-medication”. The classification of Y5_HY0-2 was given to any individual who was never classified by any rater as above HY stage 2. We also measured PIGD using the PIGD score derived by Jankovic et al at each follow up visit.^9,10^ Average year 5 PIGD scores were also derived from “on-medication” evaluations in years 4.5-6 typically generated over 3 visits, to generate a year 5 outcome variable (Y5_PIGD). Note the HY- based classification neatly dichotomizes our sample into two groups that, during treatment (“on-medication”), are either with, or without substantial progression of gait dysfunction, over the first 5 years of disease (Figure 1, “Change in PIGD Score Over Time”).

**Figure 1:**
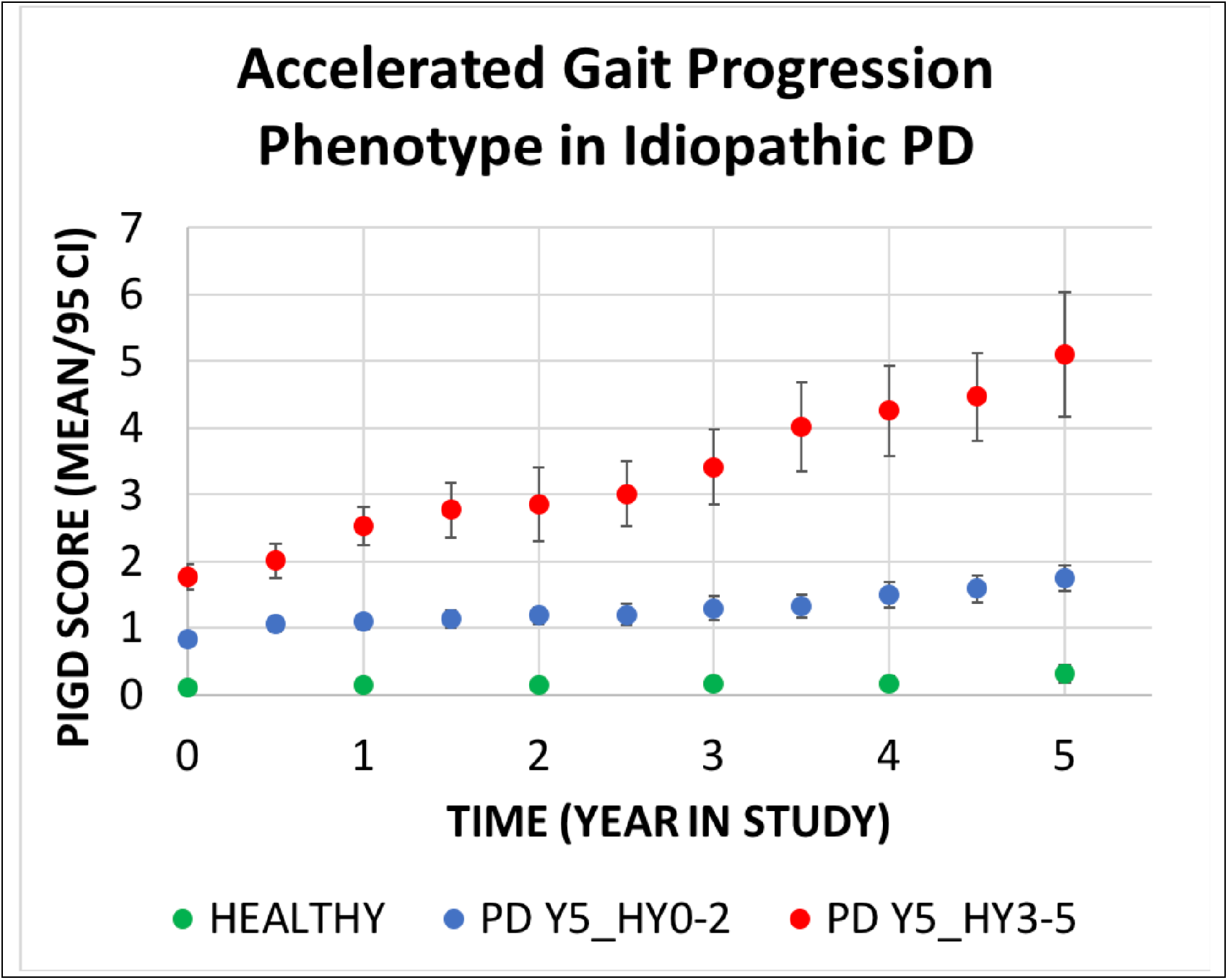
Individuals who had at least one rating of HY 3, on medication, within the first 5 years of PPMI follow up were characterized by significantly more rapid progression of gait dysfunction as measured by the PIGD scale (red circles) compared to individuals who maintained HY status of 2 or less during the first 5 years of treatment (blue circles). Controls (green) are presented as a reference.

For **<u>Scale Validation</u>**, we performed a survival analysis to evaluate the relationship between scale value (quartile rank) and population time to progression of gait related disability (at least one practitioner evaluation of HY ≥ 3) in both validation sets, using a Cox proportional hazard regression model. An additional analysis compares the utility of a 5-factor model (age, gender, and UPDRS items I, II, and III) compared to a 3-factor model (age, gender, and eGDSS) to predict progression (Y5_PIGD).

## RESULTS

### Sample Characteristics

In the de-novo cohorts, the PD sample was similar to the heathy control sample. PD subjects developing Y5_HY3-5 were older and had more severe signs and symptoms at entry (Table 2), and were characterized by progressive gait dysfunction over time as measured by PIGD rating (Figure 1).

**Table 2:**
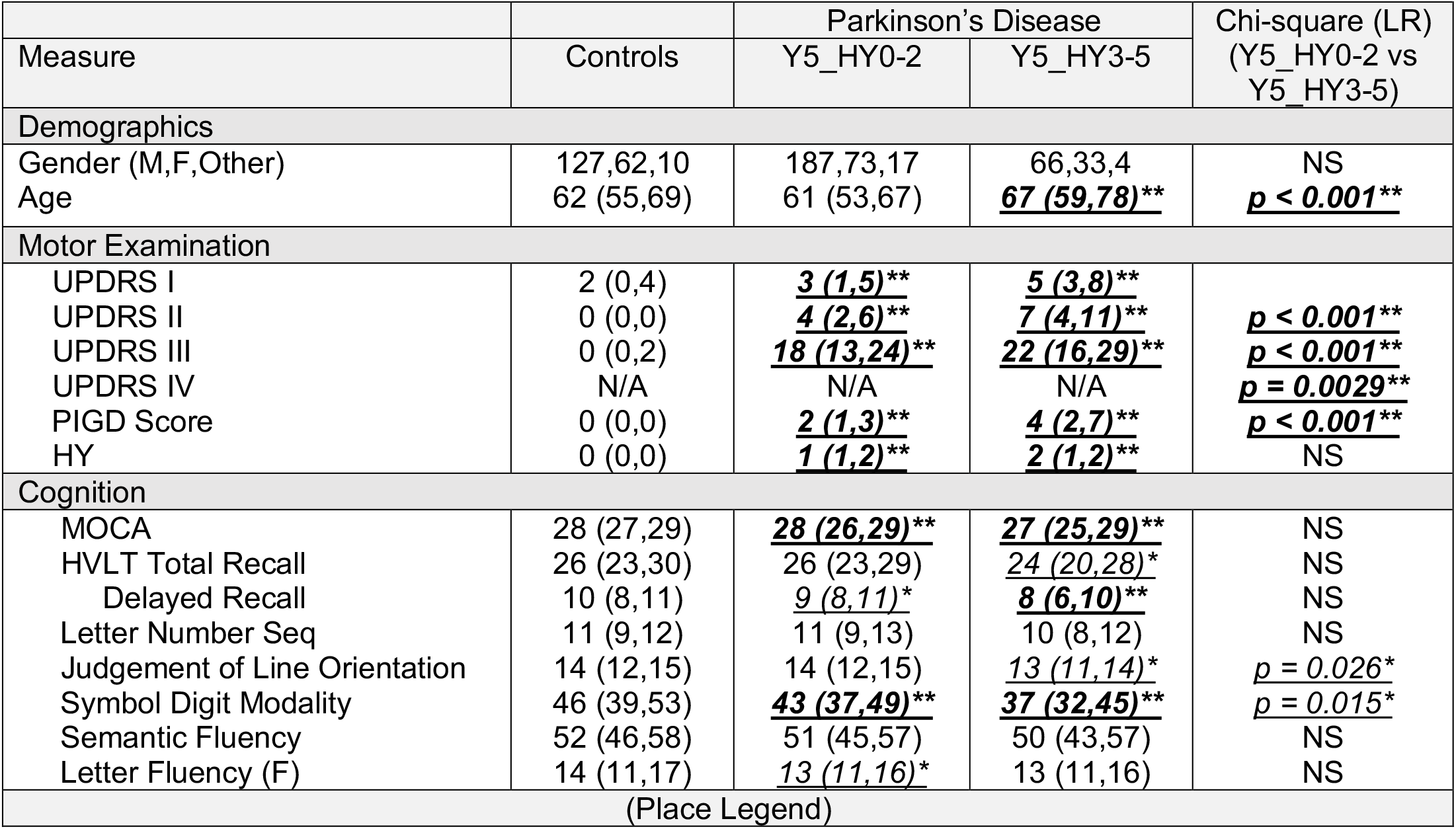
Demographics of Sample. Raw scores are presented. All evaluations have been adjusted for age, gender, and multiple comparisons. Differences between individuals who later develop progressive gait dysfunction (HY stage 3 or higher by 5 years of follow up) and those who maintain stable gait function under management are reported on the far right. Differences in each category from controls within PD columns is denoted by underline and italics. * p < 0.05 (Adjusted) ** p < 0.005 (Adjusted)

### Development of Emerging Gait Dysfunction Screening Scale (eGDSS) Factors

The scale is designed to capture items in the UPDRS associated with disability (onset of postural instability, HY 3 or greater). Specifically, a logistic regression was performed comparing individual elements on the Unified Parkinson’s Disease Rating Scale (sections 1 through 3) to HY category in a derivation set of 301 individuals with de-novo PD (216 PD Y5_HY0-2 vs 85 PD Y5_HY3-5). The sample involved 53 multiple comparisons. Significant factors were complaints of lightheadedness and fatigue (from UPDRS I), self-perception of alterations in speech, walking, and ability to rise (from UPDRS II), and objective findings of difficulty rising from a seated posture and visible stooping (from UPDRS III) at baseline, which met our criteria of p < 0.01 (adjusted for multiple comparisons) as predictors of more rapid progression to HY stage 3 or higher within 5 years of disease diagnosis (Table 3).

**Table 3:** Factors Associated with Progression to HY Stage 3 or Worse Within The First 5 Years of Disease. After correction for multiple comparisons (53 items were compared), 7 baseline UPDRS items were associated at p < 0.01 with Y5_HY3-5, and were used to develop an early gait dysfunction screening scale (eGDSS). Two other items (difficulty turning in bed and adjusting clothes, from UPDRS section II, and pain and other sensations from UPDRS I), were significant at p < 0.05 after multiple comparisons correction, but were not included. Weights were derived from a logistical regression of items (present or absent) vs. status (PD Y5_HY3-5 vs PD Y5_HY0-2).

| Item (UPDRS Section) | PD Y5_HY0-2<br>Median<br>(1<sup>st</sup> Q/3<sup>rd</sup> Q) | PD<br>Y5_HY3-5<br>Median<br>(1<sup>st</sup> Q/3<sup>rd</sup> Q) | Significance<br>(Univariate) | Multivariate<br>Logistical Weight<br>(Rounded<br>Multiple of Min) | Proposed<br>Weight |  |
| --- | --- | --- | --- | --- | --- | --- |
|  |  |  |  |  | ***Present*** | ***Absent*** |
| Lightheadedness on Standing (I) | 0 (0,0) | 0 (0,1) | ***p = 0.0042*** | ***0.91 (4)*** | ***4*** | ***0*** |
| Fatigue (I) | 0 (0,1) | 1 (0,1) | ***p = 0.0060*** | ***0.21 (1)*** | ***1*** | ***0*** |
| Speech (II) | 0 (0,0) | 0 (0,1) | ***p = 0.0023*** | ***0.85 (4)*** | ***4*** | ***0*** |
| Getting out of bed, car, or deep chair (II) | 0 (0,1) | 1 (0,1) | ***p = 0.00173*** | ***0.21 (1)*** | ***1*** | ***0*** |
| Walking and Balance (II) | 0 (0,0.25) | 1 (0,1) | ***p = 0.00051*** | ***0.74 (4)*** | ***4*** | ***0*** |
| Arising from chair (III) | 0 (0,0) | 0 (0,1) | ***p = 0.0055*** | ***1.22 (6)*** | ***6*** | ***0*** |
| Posture (III) | 0 (0,1) | 1 (0,1) | ***p = 0.0059*** | ***0.46 (2)*** | ***2*** | ***0*** |
| (Place Legend) |  |  |  |  |  |  |

### Development of eGDSS Weighting (Table 3)

Median and quartile analysis revealed presence (1) or absence (0) of the factor was a dominant characteristic at baseline. Specifically, median score, bottom quartile (25 percentile) score, and top quartile (75 percentile) score reveal scores universally between 0 and 1 at the quartiles. Therefore, to improve scale simplicity all items were binarized to present or absent, and a second logistic regression was performed, including all factors originally identified. Scale weighting for each factor was created by dividing individual weights of each item by the minimum weight of the lowest weighted item, and rounding. Table 3 shows statistical properties of the group comparison, median and quartile properties by group, and weighting strategy.

### Properties of the eGDSS Scale (Table 4, Figure 2)

Properties of the eGDSS in the derivation sample (all 380 de-novo subjects) and both validation samples shows that all individuals with PD, regardless of quartile, were (as expected) more likely to develop postural instability than controls – but quartile rank (and most specifically membership in quartile 4) was a substantial additional predictor. Table 4 shows Cox Proportional Hazard Ratio for development of disability by eGDSS Quartile in the full sample (N=380), in the IPD validation set (N=79), and in the GPD validation set (N=141). Kaplan Meier Survival Curves are shown in Figure 2 in the derivation sample (Top), the IPD de-novo validation sample (Middle), and the GPD validation sample (Bottom). Quartile 4 (eGDSS ≥ 11) is associated in both the derivation, and validation set with a highly significant increased likelihood of developing HY stage ≥ 3. The eGD phenotype will be most effective if it is a prognostic indicator for development of postural instability. We therefore evaluate the temporal relationship of the paired relationship between development of the eGD phenotype (eGDSS Score ≥ 11) and development of postural instability (HY ≥ 3) in both the IPD and the GPD cohorts. Any instance of either eGDSS Score ≥ 11, or HY ≥ 3, was considered a potential paired eGD-HY “event”, in the full sample of 380 individuals we therefore had 301 eGD “events” (79 individuals had neither occurrence). In the IPD cohort, eGDSS Score ≥ 11 occurred prior to HY ≥ 3, or HY ≥ 3 had not yet occurred, in 289 of 301 pairings, and HY ≥ 3 occurred before eGDSS ≥ 11 (or eGDSS ≥ 11 had not occurred at time of end of follow up) in 12 cases; (Chi-Square Statistic = 510, p < 0.00000001). Median time from eGDSS Score ≥ 11 to HY ≥ 3, in those who developed postural instability, was 983 days (1^st^ Quartile 366, 4^th^ Quartile 1522).

**Table 4:** Baseline eGDSS Quartile Ranking Predicts Likelihood of Advancing HY Status. Progression to HY ≥ 3, by quartile, compared to healthy controls (right-most columns) and compared to Quartile I. Reference Quartile I values in all cases are taken from the de-novo sample. A: Significant, and progressive impact of quartile on risk development of HY ≥ 3 is noted in the de-novo PD derivation sample. B: In the IPD validation sample, eGDSS scores in the fourth quartile (≥ 11) are associated with a significantly higher likelihood of progression to HY ≥ 3. C: In the GPD cohort, once again membership in Quartile 4 confers a higher risk of progression to HY ≥ 3 compared to de-novo subjects with eGDSS scores ≤ 2. ** p ≤ 0.01 *** p ≤ 0.005 **** p ≤ 0.0001

|  | eGDSS Quartile | Number in Quartile | Quartile Value (Score, Baseline) | Within Group Hazard Ratio (HR) Compared to PD Quartile I (95% CI for HR) | z | Pr (>\|z\|) | Hazard Ratio (HR) Compared to Controls (95% CI for HR) |
| --- | --- | --- | --- | --- | --- | --- | --- |
|  | Controls | 186 | Median = 2 | Control Sample |  |  |  |
| A: Derivation Sample (N=380) | I | 111 | ≤ 2 | Derivation Reference Group |  |  | 5.3 (1.7-16.3)*** |
|  | II | 85 | 3-6 | 2.03 (1.00-4.11) | 1.97 | 0.049 | 10.7 (3.7-31.5)**** |
|  | <u>III</u> | <u>78</u> | <u>7-10</u> | <u>3.34 (1.71-6.54)***</u> | <u>3.52</u> | <u>0.00043</u> | 17.7 (6.2-51.0)**** |
|  | <u>IV</u> | <u>76</u> | <u>≥ 11</u> | <u>8.28 (4.48-15.29)****</u> | <u>6.75</u> | <u>< 0.00001</u> | 44.1 (15.9-122.2)**** |
| B: IPD Sample (N=79) | I (Ref) | 86 | Derivation Sample De-Novo Quartile I Reference Group (Modified) |  |  |  |  |
|  | I | 25 | ≤ 2 | 1.90 (0.51-7.02) | 0.96 | 0.34 | 9.7 (1.9-48.4)** |
|  | II | 18 | 3-6 | 2.52 (0.68-9.29) | 1.38 | 0.17 | 12.7 (2.6-63.6)*** |
|  | III | 21 | 7-10 | 2.53 (0.68-9.47) | 1.38 | 0.17 | 13.2 (2.6-67.3)*** |
|  | <u>IV</u> | <u>14</u> | <u>≥ 11</u> | <u>16.18 (6.34-41.23)****</u> | <u>5.83</u> | <u>< 0.00001</u> | 88.0 (23.2-333.5)**** |
| C: GPD Sample (N=141) | I (Ref) | 111 | Derivation Sample De-Novo Quartile I Reference Group |  |  |  | 5.3 (1.7-16.3)*** |
|  | I | 20 | ≤ 2 | 2.81 (0.56-14.06) | 1.26 | 0.21 | 21.6 (2.5-185.3)*** |
|  | II | 29 | 3-6 | 0.94 (0.11-7.87) | -0.056 | 0.96 | 7.4 (0.6-95.1) |
|  | III | 43 | 7-10 | 3.84 (1.15-12.83) | 2.19 | 0.029 | 28.5 (4.7-174.1)*** |
|  | <u>IV</u> | <u>60</u> | <u>≥ 11</u> | <u>13.26 (5.13-34.28)****</u> | <u>5.34</u> | <u>< 0.00001</u> | 103.2 (18.7-569.9)**** |
| (Place Legend) |  |  |  |  |  |  |  |

**Figure 2:**
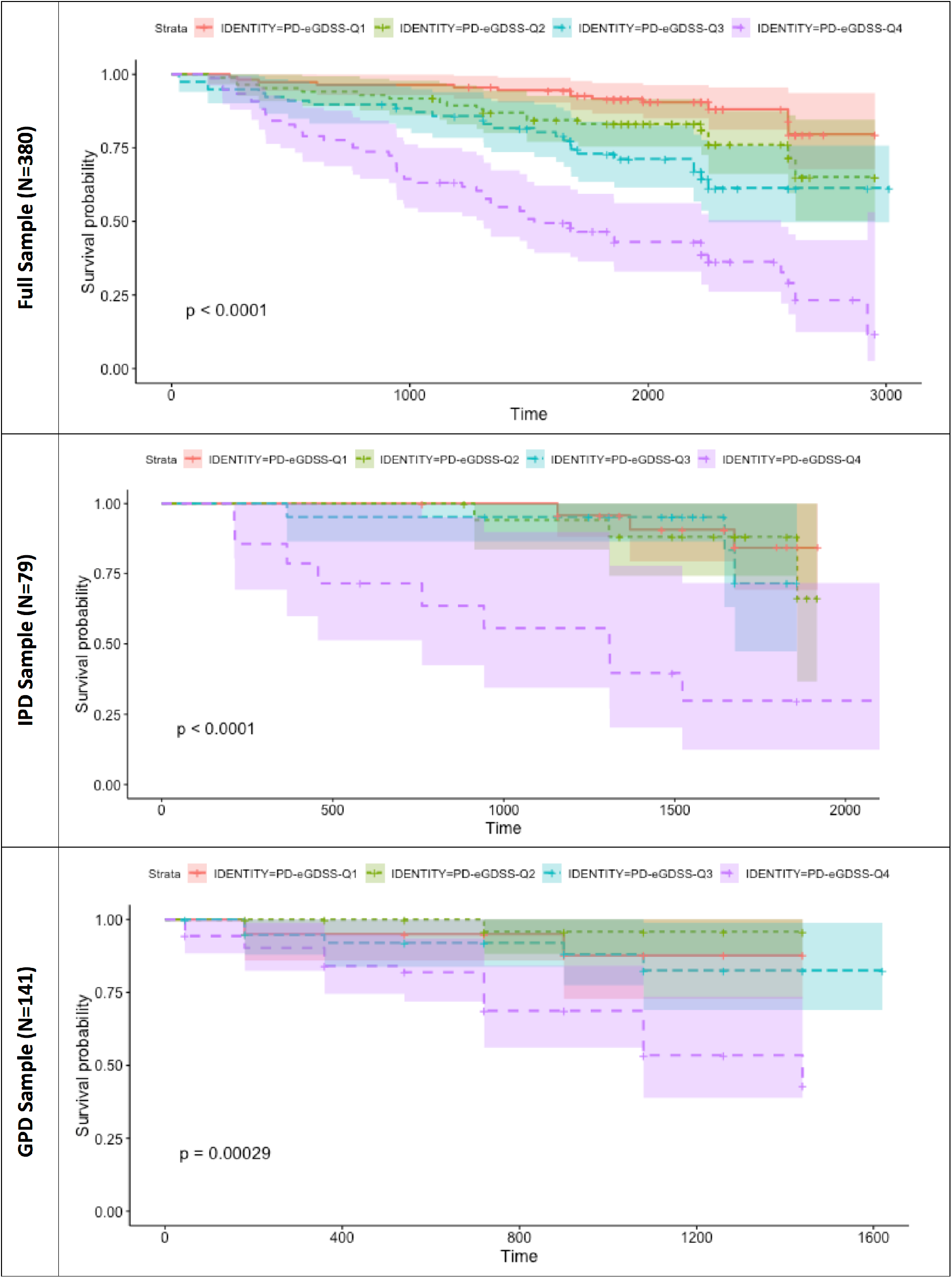
Properties of the eGDSS are shown in the full sample (Top) and in two validation sets. In both validation sets, membership in eGDSS quartile 4 is a significant predictor of later development of postural instability (HY ≥ 3) compared to both controls, and membership in Quartile I.

### Relationship of eGDSS score to cognitive decline in PD (Table 5)

As we observe eGDSS Quartile 4 substantially deviates from other quartiles in both derivation and validation analyses, we evaluated relationship of membership in Quartile 4 and slope of cognitive change, compare eGDSS Quartile 4 to eGDSS Quartile 1. Membership in Quartile 4 conferred a significant increased likelihood of cognitive progression, as measured by the MOCA and HVLT, after correction for multiple comparisons. Rate of change in UPDRS in de-novo subjects is not different in relation to eGDSS quartile, and rate of cognitive change does not differ significantly between Quartiles 1, 2, and 3 (not shown). A repeat analysis including Quartiles 1-3 versus Quartile 4 resulted in similar results to those shown in Table 5 (not shown).

**Table 5:**
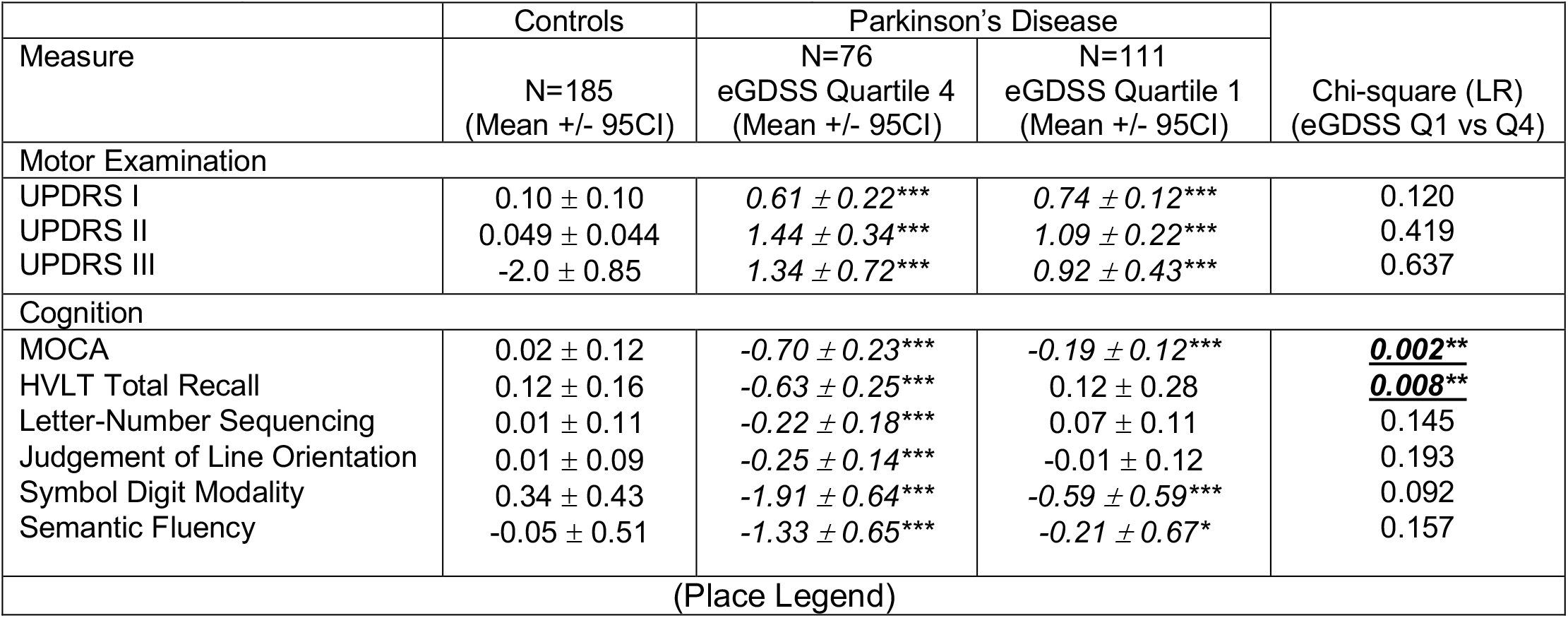
Change in Motor, Neurobehavioral, and Cognitive Function in Sample. Slope (mean point change per year), with 95% confidence intervals, for motor and cognitive scores in PPMI for all de-novo subjects with at least 5 years of data available. Significance is adjusted for age and gender; for cognitive measures significance of 0.00833 (0.05/6) is to achieve statistical threshold. * p < 0.05 ** p < 0.00833 *** p < 0.0001

| Table 5: Change in Motor, Neurobehavioral, and Cognitive Function in Sample |  |  |  |  |
| --- | --- | --- | --- | --- |
| Measure | Controls | Parkinson's Disease |  | Chi-square (LR)<br>(eGDSS Q1 vs Q4) |
|  | N=185<br>(Mean +/- 95CI) | N=76<br>eGDSS Quartile 4<br>(Mean +/- 95CI) | N=111<br>eGDSS Quartile 1<br>(Mean +/- 95CI) |  |
| Motor Examination |  |  |  |  |
| UPDRS I | 0.10 ± 0.10 | 0.61 ± 0.22*** | 0.74 ± 0.12*** | 0.120 |
| UPDRS II | 0.049 ± 0.044 | 1.44 ± 0.34*** | 1.09 ± 0.22*** | 0.419 |
| UPDRS III | -2.0 ± 0.85 | 1.34 ± 0.72*** | 0.92 ± 0.43*** | 0.637 |
| Cognition |  |  |  |  |
| MOCA | 0.02 ± 0.12 | -0.70 ± 0.23*** | -0.19 ± 0.12*** | ***0.002\*\**** |
| HVLT Total Recall | 0.12 ± 0.16 | -0.63 ± 0.25*** | 0.12 ± 0.28 | ***0.008\*\**** |
| Letter-Number Sequencing | 0.01 ± 0.11 | -0.22 ± 0.18*** | 0.07 ± 0.11 | 0.145 |
| Judgement of Line Orientation | 0.01 ± 0.09 | -0.25 ± 0.14*** | -0.01 ± 0.12 | 0.193 |
| Symbol Digit Modality | 0.34 ± 0.43 | -1.91 ± 0.64*** | -0.59 ± 0.59*** | 0.092 |
| Semantic Fluency | -0.05 ± 0.51 | -1.33 ± 0.65*** | -0.21 ± 0.67* | 0.157 |
| (Place Legend) |  |  |  |  |

### Relationship of eGDSS score to Y5_PIGD (Table 6)

While the eGDSS scale is keyed to disability as measured by the HY scale, the relationship between eGDSS and development of the broader PIGD phenotype is of interest. While age, gender, and UPDRS 1, 2, and 3 (5 factors) accounted for only 19.9% of variance in predicting Y5-PIGD, the proposed eGDSS scale (3 factors) accounted for an adjusted 26.1% of variance (p = 0.018 – see table 6). In summary, eGDSS score at baseline explained more variance in Y5_PIGD than UPDRS, and UPDRS did not add significantly to the eGDSS score with regards to explaining variance in Y5_PIGD.

**Table 6:** Relative Relationship eGD Phenotype to PIGD Phenotype. While age is superior to the UPDRS in predicting progression, the baseline eGDSS score is superior to age in the de-novo validation set (N=79). ** p = 0.015

| Table 6: Relative Relationship eGD Phenotype to PIGD Phenotype |  |  |  |  |  |
| --- | --- | --- | --- | --- | --- |
| SOURCE | F | Pr > F | SOURCE | F | Pr > F |
| AGE | 8.282 | 0.005 | **eGDSS** | **15.999** | <0.00001 |
| UPDRS I | 0.753 | 0.388 | AGE | 5.354 | 0.023 |
| UPDRS II | 2.040 | 0.158 | GENDER | 0.000 | 0.998 |
| UPDRS III | 3.059 | 0.085 |  |  |  |
| GENDER | 0.177 | 0.675 |  |  |  |
| (ADJUSTED)<br>EXPLAINED VARIANCE |  | 0.199 | (ADJUSTED) EXPLAINED<br>VARIANCE |  | 0.274** |

### Post-hoc analyses – capacity of baseline PIGD scale to predict Y5_PIGD and Y5_HY3-5

A pressing question when developing a new metric is whether it improves on existing metrics. We therefore evaluated if baseline PIGD provides relevant information to predict later HY status and PIGD scores. Baseline PIGD was a poor predictor of Y5_PIGD in both the derivation (3 factors, adjusted R^2^ = 0.172) and de-novo validation set (adjusted R^2^ = 0.169), and did not improve prediction of Y5_PIGD when added to the UPDRS (R^2^ = 0.254, similar to UPDRS alone). In a 4 factor ANOVA model, baseline PIGD (F statistic = 0.011, Pr > F = 0.917), did not improve prediction when added to the proposed eGDSS scale, and adjusted estimated accounted variance (25.1%) declined due to addition of a factor. In a logistic model (prediction of Y5_HY3-5), adding PIGD to eGDSS (4 factors, area under ROC curve = 0.784) did not improve prediction of the 3-factor model (Age, Gender, and eGDSS alone, area under ROC curve = 0.784).

## DISCUSSION

Specific clinical features, years prior to the onset of disabling gait dysfunction, predict development of postural instability in PD. Based on our findings, we propose an emerging gait disorder phenotype (eGD) can be detected at baseline, and that individuals who display features of this phenotype are at high risk of accelerated development of both gait and cognitive dysfunction. We operationalize detection of the eGD phenotype using a provisional emerging gait dysfunction phenotype screening scale (eGDSS). While more detailed studies to develop ideal cutoff scores would be appropriate, at this stage and our data suggest that an eGDSS score ≥ 11 is a reasonable criteria for identifying the phenotype. Individuals in this quartile had substantially increased risk of developing disabling postural instability as measured by the HY rating scale, and showed accelerated cognitive progression as measured by the MOCA and HVLT. The baseline eGDSS score also explained more variance in PIGD score in year 5 than the UPDRS. The proposed eGD phenotype showed itself to be robust in two separate validation sets: (1) a separate de-novo set of individuals with idiopathic PD and (2) a validation set of individuals with variable time of onset of symptoms with one of three known genetic causes of PD. Finally, the proposed eGD phenotype occurs before development of postural instability as measured by the HY scale (by a median of nearly 3 years), indicating the phenotype is potentially predictive of deterioration, rather than simply an associated phenotype. The eGDSS scale has an acknowledged weakness in relying significantly on patient self-reports, and we do not at this stage attempt to evaluate clear cut-off points for phenotypic identification. This initial version of the eGDSS scale may therefore be useful in and of itself, but we consider it most useful at this stage to provide guidance in developing an objective baseline evaluation to more clearly define the phenotype.

The clinical features we identify in this work suggest some avenues for further research on understanding the pathophysiology of the gait dysfunction in PD. Specifically, we find the following features appear to be associated with more rapid progression of gait dysfunction: (1) lightheadedness on standing, (2) self-perceived speech difficulties, (3) difficulties standing/rising, (4) postural changes, and (5) visible changes in the substantia nigra, as well as changes in dopaminergic projects as measured by DATscan.^29^ A lesion in the substantia nigra could be associated with some of these clinical findings, including changes in posture and postural reflexes, and speech changes such as hypophonia, however our assessment does not clearly define the source of self-perceived language deficits in this population, which could also be related to a lesion in the dorsal motor nucleus of the vagus (one of the first regions impacted by lewy bodies in PD in Braak’s formulation),^30-32^ or might even represent a cognitively-based symptom. Complaints of lightheadedness on standing suggests either a peripheral or central (brainstem) lesion in noradrenergic and adrenergic neurons. The association of these apparently diffusely distributed brainstem-related symptoms with more rapid cognitive deterioration suggests a more aggressive disease process.

In summary, we show that risk for developing postural instability, and the PIGD phenotype, is detectable at baseline clinically, based on symptoms and clinical findings (the eGD phenotype) that are distinct from those that later characterize the PIGD phenotype. Identifying a sub-population at baseline at high risk of clinical progression is important from the perspective of both understanding the pathophysiology of PD progression, and for identifying individuals in which disease modifying agents can be evaluated. From a practical perspective, a rapid clinical evaluation that can define future risk of disability is particularly useful. Heretofore, the Unified Parkinson’s Disease Rating Scale (UPDRS) has been a primary focus and target in intervention studies. However summary UPDRS metrics are primarily useful within subject for comparing the impact of medications on symptoms and motor findings of disease, and are less useful for evaluating the impact of potential disease-modifying therapies.^6^ We show here that for the majority of individuals in the first 5 years of observation in PPMI (the first 6 years of diagnosed disease), changes in gait function under management are modest, but progressive in a sub-population despite treatment. By formally identifying this eGD phenotype we identify a population group destined to develop dopamine resistant symptoms that can targeted for early intervention studies.

## Acknowledgement statement (including conflict of interest and funding sources)

There are no conflicts of interest associated with this publication. We would like to acknowledge the funding of the National Institutes of Health through ***K23NS083620***, and grant funding from the Michael J Fox Foundation in support of this research. Data used in the preparation of this article were obtained from the Parkinson’s Progression Markers Initiative (PPMI) database (www.ppmi-info.org/data). For up-to-date information on the study, visit www.ppmi-info.org. PPMI – a public-private partnership – is funded by the Michael J. Fox Foundation for Parkinson’s Research and funding partners, including Abbvie, Allergan, Avid Radiopharmaceuticals, Biogen, Biolegend, Bristo-Myers Squib, Denali, GE Healthcare, Genentech, GlaxoSmithKline, Lilly, Lundbeck, Merck, Meso Scale Discovery, Pfizer, Roche, Sanofi Genzyme, Servier, Takeda, Teva, and UCB (see www.ppmi-info.org/fundingpartners).

There are no conflicts of interest associated with this publication. We would like to acknowledge the funding of the National Institutes of Health through ***K23NS083620***, and grant funding from the Michael J Fox Foundation in support of this research. Data for this research was obtained from the Parkinson’s Progressive Markers Initiative.

## Author Contributions

FMS, ED, and DS participated in study concept and design. FMS, WSM, CH, TA, AG, LJ, AB, TD, and GC participated in data acquisition and analysis. FMS, ED, WSM, and CH participated in drafting the manuscript and constructing figures.

